# DNA methylation modules associate with incident cardiovascular disease and cumulative risk factor exposure

**DOI:** 10.1101/471722

**Authors:** Kenneth Westerman, Paola Sebastiani, Paul Jacques, Simin Liu, Dawn DeMeo, José M. Ordovás

**Affiliations:** JM-USDA Human Nutrition Research Center on Aging at Tufts University, Boston, MA, USA; Department of Biostatistics, Boston University School of Public Health, Boston, MA, USA; Department of Epidemiology, Brown University, Providence, RI, USA; Channing Division of Network Medicine, Department of Medicine, Brigham and Women’s Hospital, Boston, MA, USA; IMDEA Alimentación, CEI, UAM, Madrid, Spain; Centro Nacional de Investigaciones Cardiovasculares (CNIC), Madrid, Spain

## Abstract

**Abstract:** *Background:* Epigenome-wide association studies using DNA methylation have the potential to uncover novel biomarkers and mechanisms of cardiovascular disease (CVD) risk. However, the direction of causation for these associations is not always clear, and investigations to-date have generally failed to replicate at the level of individual loci. Here, we undertook module- and region-based DNA methylation analyses of incident CVD in the Women’s Health Initiative (WHI) and Framingham Heart Study Offspring Cohort (FHS) in order to find more robust epigenetic biomarkers for cardiovascular risk.

*Methods and Findings:* We applied weighted gene correlation network analysis (WGCNA) and the Comb-p algorithm to find methylation modules and regions associated with incident CVD in the WHI dataset. We discovered two modules whose activation correlated with CVD risk and replicated across cohorts. One of these modules was enriched for development-related processes and overlaps strongly with epigenetic aging sites. For the other, we showed preliminary evidence for monocyte-specific effects and statistical links to cumulative exposure to traditional cardiovascular risk factors. Additionally, we found three regions (associated with the genes SLC9A1, SLC1A5, and TNRC6C) whose methylation associates with CVD risk.

*Conclusions:* In sum, we present several epigenetic associations with incident CVD that reveal disease mechanisms related to development and monocyte biology. Furthermore, we show that epigenetic modules may act as a molecular readout of cumulative cardiovascular risk factor exposure, with implications for the improvement of clinical risk prediction.

## 2 Introduction

Genetic approaches to cardiovascular disease (CVD) research have led to important breakthroughs in mechanistic understanding and therapeutic strategies. However, the mechanisms for gene variant-disease relationships are often difficult to determine, and their effects may often be mediated by epigenetic regulation [1]. DNA methylation is one such mechanism that can reflect both genetic variation and environmental exposures and potentially drive their effects on CVD outcomes [2].

A series of recent epigenome-wide association studies (EWAS) have examined relationships between DNA methylation at cytosine-phosphate-guanine (CpG) sites and various subtypes of CVD, including prior myocardial infarction [3], acute coronary syndrome [4], and atherosclerosis [5]. These cross-sectional studies may reveal important mechanistic insights, but are susceptible to reverse causation, i.e. methylation being influenced by the presence of CVD. Indeed, Mendelian randomization approaches across multiple phenotypes have suggested that reverse causation is more common [6,7] than the causal methylation effect that is often implicitly assumed. One approach to this problem is to examine epigenetic associations with cardiovascular risk factors. Multiple investigations have explored these relationships genome-wide [8,9], and have even uncovered prognostic CpG sites for incident coronary heart disease in the process [10,11]. A few studies looking directly at incident CVD as a binary variable have found relationships with global DNA methylation (as approximated by LINE-1 methylation levels) and with a specific cluster of CpG sites in the ZBTB12 gene [12,13].

Studies linking CVD and methylation have additionally shown a notable lack of replication, especially at the level of single CpG sites [14]. One approach to this problem is to aggregate CpGs and test their phenotype associations at the group level. Differentially methylated region (DMR) searches may improve detection by combining sites based on physical proximity on the genome [15,16]. An alternative grouping strategy is to search for correlation-based clusters, which may boost biological signal and improve the interpretability of results [17]. This approach was originally developed for use with gene expression data, but has been successfully applied to higher-dimensional DNA methylation microarray datasets [18,19].

To address the problem of reverse causation by CVD while achieving more robust results, we set out to analyze relationships between group-level CpG methylation and incident CVD using time-to-event models in two cohorts. We used module- and region-based techniques to improve detection and provide more interpretable results. We sought context for two specific modules of interest using gene- and chromatin-based annotations, and compared module activations to past and current cardiovascular risk factor levels to better understand their potential biological mechanisms.

## 3 Results

### 3.1 Weighted correlation network approach finds CVD-related modules

Population characteristics are described in Table 1. The discovery set, Women’s Health Initiative (n=2023), had a median age of 65 at blood draw and is entirely female, while being selected for an approximately equal ratio of subjects who did and did not experience an incident CVD event following the methylation measurement timepoint. The replication set, Framingham Heart Study Offspring Cohort (n=2587), had a median age of 66 at blood draw (Exam 8) and is approximately half female, with 305 subjects experiencing incident CVD events. Cardiovascular events were defined here as encompassing coronary heart disease, stroke, and death from CVD (see Methods section for further details).

**Table 1:** Population description

|  | WHI | FHS |
| --- | --- | --- |
| Sample Size | 2023 | 2587 |
| % female | 100 % | 55 % |
| Age at blood draw | 65 (59-70) | 66 (60-73) |
| Mixed race | Yes | No |
| BMI | 29.1 (25.5-33.3) | 27.7 (24.5-31) |
| % smoke currently | 10% | 9% |
| Smoking pack-years | 0 (0-12.5) | 0 (0-0) |
| # prior CVD events | 0 | 331 |
| # incident CVD events | 1009 | 305 |

<sup>1</sup> Continuous values shown as: median (interquartile range).
<sup>2</sup> 92 subjects experienced both prior and incident CVD events.

We first set out to find biologically relevant modules in an unsupervised manner (agnostic to incident CVD information) using the WGCNA algorithm for 422952 CpGs in WHI passing quality control filters (study overview in Supp. Fig. S1). After weighted correlation network construction, topological overlap calculation, and subsequent clustering, 110 modules were uncovered, ranging in size from 28 to 35361 CpGs.

Principal component eigenvectors for each module were calculated in order to examine the characteristics of these modules as a whole. The first principal component of each module tended to explain approximately half of the total variance, while the rest contributed only small fractions (see Supp. Fig. S2 for selected Scree plots). Thus, these first eigenvectors, or “eigenCpGs”, were subsequently used to describe module behavior. Cox proportional hazards models were used to assess the relationships between these module eigenCpGs and incident CVD. In partially-adjusted models (adjusted for technical factors and estimated white blood cell proportions), three modules were found to be associated at multiple test-corrected FDR < 0.2 (Table 2; correction based on 110 modules). Adjustment for biological covariates (age, BMI, sex/race, and smoking behavior) attenuated these relationships to marginal statistical significance (all 0.01 < p < 0.1; direct risk factor associations shown in Fig. 3). These modules showed strong (FDR < 10^-4^) enrichment for different sets of GO terms, ranging from immune activation (myeloid or T cell) to developmental processes.

**Table 2:** Modules associated with incident CVD at FDR < 0.2

| module | size | EigenCpG |  | Enrichment analysis |  |  |
| --- | --- | --- | --- | --- | --- | --- |
|  |  | Var. expl. (%) | p | GO terms | CpG Islands | Gene-based |
| blue | 29441 | 44.61 | 0.00027 | Development | N_shore | 1stExon/TSS/5' UTR |
| brown | 953 | 53.08 | 0.00455 | Immune activation | Open sea |  |
| purple | 568 | 44.88 | 0.00500 | T cell activation | Open sea | Body |

All three modules showed very strong preservation in FHS (all *Z_summary_* statistics > 50, where 10 is a typical threshold for strong preservation), when evaluated using established density and connectivity preservation techniques [20]. Of these, two associations with incident CVD (blue and brown) replicated strongly in FHS, while purple showed nominal replication (p=0.0203) in partially-adjusted models (Supp. Table 1). Fully-adjusted models including age as a covariate attenuated (brown) or abolished (blue and purple) these associations in FHS.

Though the existence of past CVD events (experienced prior to sample collection for DNA methylation measurement) could represent a confounder in the FHS dataset, sensitivity analyses adjusting for past events did not appreciably reduce the strength of these module-trait relationships. Also of potential relevance to this replication is the demographic heterogeneity between the two cohorts. To address this possibility, we performed additional analyses including interaction terms between eigenCpGs for each module and either sex (in FHS) or race (in WHI). None of these analyses produced significant interaction terms at p < 0.05.

### 3.2 Genome-wide associations between DNA methylation and incident CVD events

To investigate more specific DNA methylation signals, we performed an epigenome-wide association study (EWAS) for incident CVD. Of single sites from the EWAS, 3 reached a genome-wide Bonferroni threshold, but none replicated strongly in FHS (Supp. Table 2). In order to improve statistical power, we focused on differentially methylated regions (DMRs) with respect to incident CVD status. Single-site EWAS p-values were used as input to the Comb-p algorithm, which seeks regions enriched for low p-values while accounting for autocorrelation based on genomic distance. Comb-p was applied separately to EWAS results from WHI and FHS.

206 DMRs were found in WHI after Sidak multiple testing correction for each DMR based on its length. Of these, 3 were both found in FHS and replicated at a Bonferroni level (Table 3; Fig. 1). These regions were annotated to two cellular transport genes (SLC9A1 and SLC1A5) and TNRC6C, which codes for a scaffolding protein involved in miRNA-mediated translational repression. Of the three WGCNA modules identified above, brown CpG sites consituted part of 2 DMRs (at SLC9A1 & SLC1A5), while a single CpG from the blue module was also a member of the SLC9A1 DMR.

**Table 3:** Comb-p regions with multiple test-corrected p<0.05 in WHI and Bonferroni p<0.05 in FHS

| Location | # CpGs | Annotated gene | Genomic region | Discovery |  | Replication |
| --- | --- | --- | --- | --- | --- | --- |
|  |  |  |  | P | Adj. P | P |
| chr1:27440462-27440721 | 3 | SLC9A1 | Body | 1.75e-08 | 2.85e-05 | 0.000103 |
| chr19:47287777-47288263 | 6 | SLC1A5 | CpG shelf near TSS | 5.91e-07 | 0.000514 | 3.8e-10 |
| chr17:76037034-76037562 | 6 | TNRC6C | CpG island in 5' UTR | 1.67e-05 | 0.013300 | 0.000189 |

**Figure 1:**
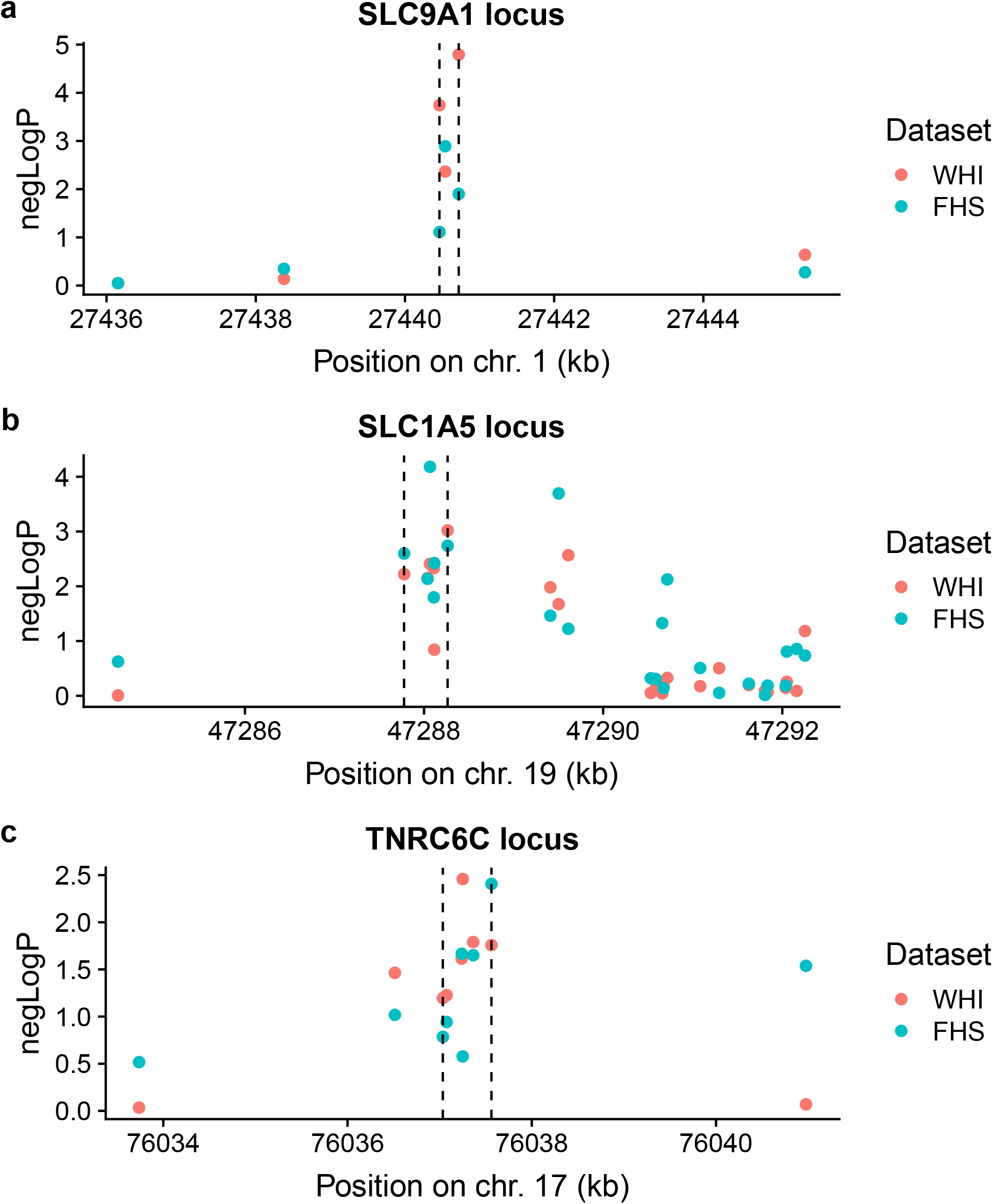
DMRs identified by Comb-p in WHI and validated in FHS at the (a) SLC9A1, (b) SLC1A5, and (c) TNRC6C loci. Negative logarithms of EWAS p-values are shown as a function of the genomic coordinate. EWAS p-values from WHI are in red and FHS in green. Dotted lines demarcate the DMR boundaries.

### 3.3 Exploration of the brown and blue modules

Based on the results from the module- and region-centric analyses, we investigated the brown and blue modules further for biological significance. The brown module was associated with immune-related genes as noted above, and was enriched strongly for “open sea” sites (p=1.1e-42) and annotated enhancers (p=1.7e-33). In contrast, the blue module was associated with development-related genes, and was enriched moderately for sites near genic transcription start sites and strongly for CpG islands (p < 2.2e-16) (Fig. 2a,b).

**Figure 2:**
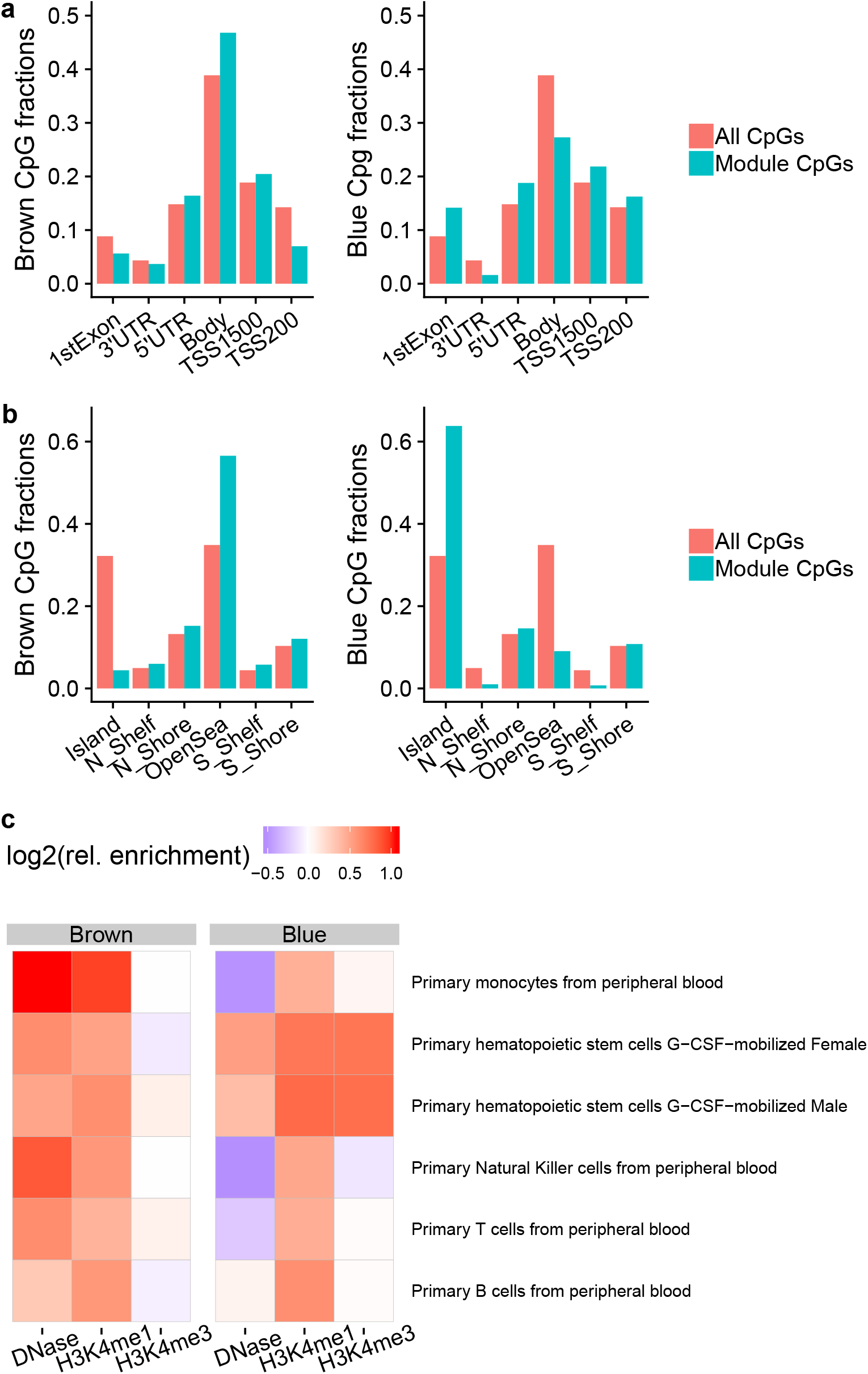
Genomic and epigenomic annotations of the brown and blue modules. a,b) Relative proportions of module CpGs compared to the full set of CpGs tested, with respect to gene-based (a) or CpG island-based (b) annotations (UTR = untranslated region; TSS_X = sites within X base pairs upstream of the gene transcription start site). c) Cell type-specific enrichments based on Roadmap Epigenomics datasets. Shown are relative enrichments of peaks (ratio of in-module fraction to all-CpG fraction) for a given epigenetic mark across many blood cell types, for each of the modules of interest.

**Figure 3:**
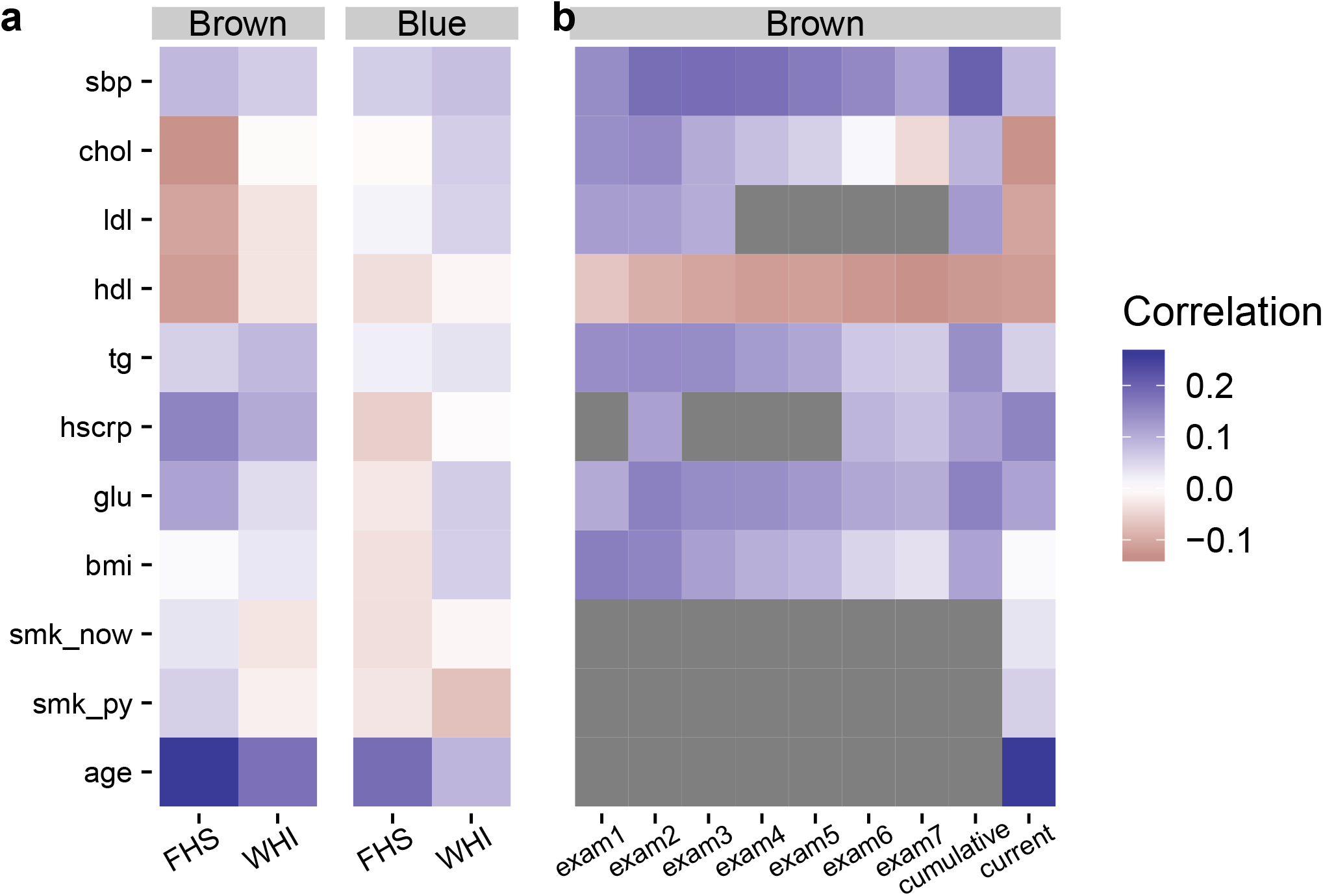
Risk factor-module relationships. (a) Pearson correlations between a series of traditional cardiovascular risk factors and module eigenCpGs (blue and brown) are shown in each study population. (b) Pearson correlations between historical risk factor levels in FHS (across previous exams, x-axis) and current brown module activation are shown. Grey panels indicate that the risk factor in question was not available for the corresponding exam.

Given these observations, we examined relative enrichments of enhancer- and promoter-associated histone marks across different blood cell subtypes to better understand the cell type specificity of this signal. Epigenetic peaks were annotated using data from the Roadmap Epigenomics Project [21] and relative enrichments were calculated as the fraction of module CpGs found in peaks divided by the fraction of all CpGs found in peaks.

We observed the greatest enrichment of brown CpGs in 2 enhancer-associated chromatin annotations, DNase hypersensitivity sites (DHS) and H3K4me1 histone peaks, from monocytes compared to other blood cell subtypes (Fig. 2c). This could point towards monocyte-related biology and inflammatory processes as an important shared mechanism for cardiovascular risk between the two cohorts examined here. In contrast, blue CpGs were enriched for DHS and promoter-associated H3K4me3 histone peaks from hematopoietic stem cells (HSCs), providing a link to the observed enrichment of develoment-related genes in this set.

The module CpG sets were also compared to two existing methylation-based age predictors from Horvath and Hannum et al., as well as the recent morbidity-directed phenoAge [22–24]. While enrichments for brown CpGs were moderate to nonexistent, blue CpGs were strongly enriched for all three of these sets, most highly for the original DNAm age developed by Horvath (46/353; p=3.4e-5; hypergeometric test), despite the fact that this model was developed based on only ~21,000 CpGs shared between multiple versions of the Illumina methylation microarray platform. Furthermore, 28 of these 46 CpGs had associated positive coefficients in the DNAm age predictor. This subset has been previously observed to contain a disproportionate amount of Polycomb-group target genes, which are known to associate with developmental processes and to be generally hypermethylated with age [25]. Using SUZ12 binding regions [26] as a proxy for Polycomb-group targets, we confirmed their enrichment in the blue module (p = 1.37e-07). Surprisingly, the blue eigenCpG showed only a modest correlation with age (r=0.09).

### 3.4 Module-risk factor relationships

Next, we examined correlations between these module eigenCpGs and traditional cardiovascular risk factors. Though no extremely strong module-risk factor correlations were observed (all |r|<0.25), they tended to be stronger for the brown module, especially in FHS (Fig. 3a). Age showed the greatest association, while lipid and glycemic parameters also showed moderate associations. To further probe relationships between the brown module and risk factors in FHS, we retrieved historical risk factors measured in previous Offspring Cohort exams. Visual inspection revealed a notably stronger correlation between the module eigenCpG and cumulative (mean of all previous exams) compared to current risk factor exposure. This pattern applied for systolic blood pressure (strongly), triglycerides, glucose, BMI, and LDL (which correlated in the “expected” direction cumulatively, but non-intuitively at Exam 8) (Fig. 3b).

To better investigate this phenomenon, we tested associations between the brown module and each of the cumulative risk factors after adjustment for potential confounders. Specifically, for each risk factor, linear models were used to predict the brown eigenCpG value from either the current or cumulative risk factor level while adjusting for the full set of EWAS covariates other than BMI (age/sex/smoking/cell counts/stu center/7 ctrl-probe PCs). Only for the brown module did cumulative risk factor exposure show strong associations, which were generally equal to or stronger than those of the current risk factors, most notably for BMI, hsCRP, and triglycerides (Table 4). Though more recent medication use could possibly explain discrepancies between biological relationships with current and past risk factors, adjustment for hypertension and lipid medication use did not notably affect the results of these models. [H]

**Table 4:**
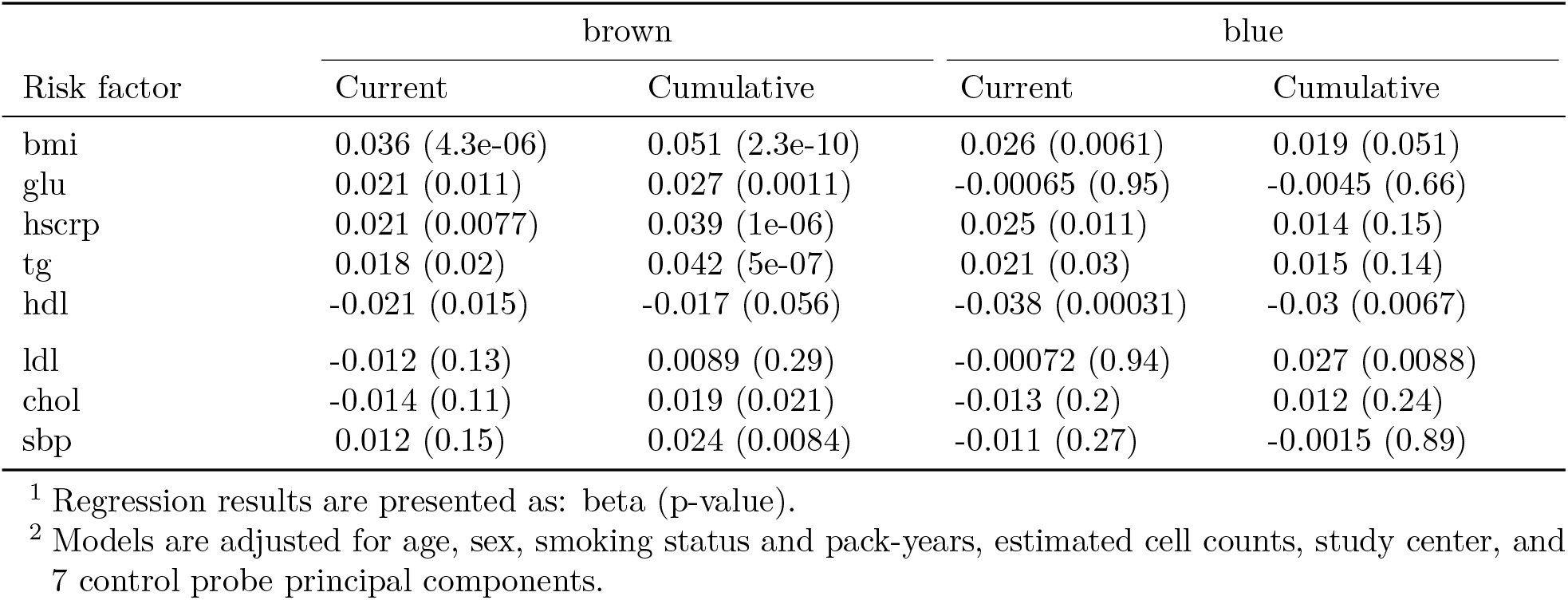
Module-risk factor relationships (current and cumulative) after adjustment for covariates

Finally, we used the basic mediation approach of Baron and Kenny [27] to test whether brown module activation may mediate a portion of the effects of cumulative risk factor exposure on cardiovascular risk. A series of Cox models were created in FHS for these three most strongly-associated risk factors (BMI, hsCRP, and triglycerides). Covariates in all models included current values for the risk factor in question, as well as technical factors, estimated cell counts, age, and sex. Current risk factors did not show notable relationships with incident CVD in any of the models. Having established the exposure-mediator relationships (Table 4), we tested the association with CVD risk of 1) cumulative risk factors, 2) module eigenCpGs, and 3) both quantities together (Table 5; example causal diagram using hsCRP in Supp. Fig. S4). In general, the significance of the module relationships with CVD tended to decrease in the presence of cumulative risk factor values. This fits with a model in which, rather than mediating cardiovascular risk, module activation acts as a biomarker for the actions of cumulative risk factor exposures by some other mechanism. As only subjects with current risk factor values were included in each model, sample sizes were largely identical across models.

**Table 5:**
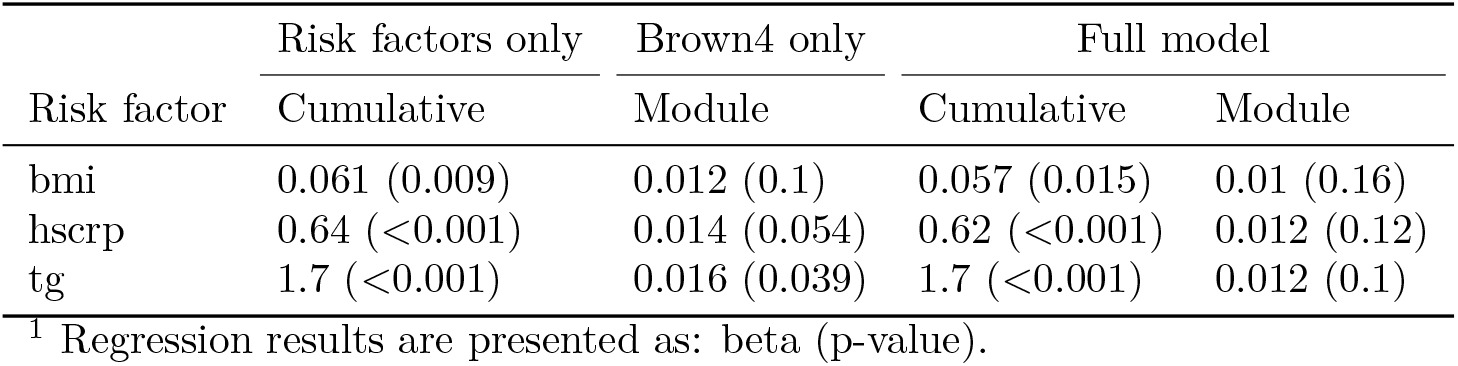
CVD risk models using cumulative risk factor exposure and brown module activation

## 4 Discussion

Here, we performed a primarily module-based epigenetic analysis of incident cardiovascular events in order to find robust, prospective biomarkers and uncover novel mechanisms contributing to disease risk. We began by constructing correlation-based clusters in the methylation data from WHI using the WGCNA algorithm. This network-based feature clustering approach can potentially improve the signal-to-noise ratio of high-dimensional DNA methylation data while facilitating more clear biological interpretation of results [28]. As WGCNA does not consider class labels (i.e. incident CVD status), the 110 modules uncovered were not *a priori* expected to be related to CVD and rather reflected unbiased patterns in the data. After correction for multiple testing, the first principal components (eigenCpGs) of three of these modules were found to be related to incident cardiovascular events. A gene ontology-based enrichment analysis of the genes annotated to these modules found strong enrichment for either immune-related or development-related processes. The finding of immune-related processes is intuitive given that DNA from blood measures primarily immune cells, while the development-related enrichment could possibly reflect influences during early life [29]. Notably, these two module “types” (immune and development) have been uncovered in a prior network-based DNA methylation analysis related to asthma [19], suggesting that similar module types are a potentially general feature of blood-based methylation patterns and that these patterns may not be fully cardiovascular-specific, reflecting instead a predisposition toward general inflammatory disease processes. Both in WHI and in replication in FHS, two modules (blue and brown) showed strong relationships with incident CVD that were attenuated after adjustment for age (direct correlations of these modules with age are presented in Fig. 3).

We examined the set of module eigenvector loadings as a proxy for the relative importance of their component CpGs, in a similar approach to the standard calculation of gene-module correlations (or “kME” statistics) in WGCNA analyses. As we did not observe any obvious peaks distinguishing particularly important groups of CpGs, we undertook an epigenome-wide association study (EWAS) in order to identify potentially stronger locus-specific signals. Though we did not find any single sites replicating in FHS after stringent correction for multiple tests, a subsequent region-based analysis using the Comb-p algorithm revealed three regions replicating strongly across the two cohorts examined here. One was found on chromosome 1 in the body of the SLC9A1 (also known as NHE-1) gene, which codes for an integral membrane ion transporter involved in intracellular pH maintenance. SLC9A1 has been shown to be required for the increased adhesion, migration, and phagocytosis of oxidized LDL seen in monocytes in response to stimuli including leptin, adrenaline, and hyperglycemia [30]. Another region discovered was on chromosome 19 near the transcription start site (TSS) of SLC1A5, which codes for a neutral amino acid transporter. Though strong evidence does not yet exist linking SLC1A5 to cardiovascular mechanisms, its CpGs have shown associations with diabetes, blood pressure, and mortality [31–33], and we note that its companion amino acid transporter, SLC7A5, is known to regulate metabolic and inflammatory reprogramming of monocytes in response to stimulation by lipopolysaccharide (LPS). Notably, CpG sites in both SLC9A1 and SLC1A5 were discovered and replicated in a recent EWAS for BMI (including the FHS cohort) [34], though the specific SLC9A1 site from that study was not one of the three constituent CpGs in the region found here. These two SLC transporter DMRs contained CpGs belonging to blue (1 in SLC9A1) and brown (1 in SLC9A1, 5 in SLC1A5) modules. The third region was found near the TSS of TNRC6C on chromosome 17. This gene codes for a component of the miRNA-mediated translational repression cascade, has shown up in a genome-wide association study (GWAS) for heart failure (not one of the phenotypes included in our CVD definition here) [35], and was identified as a potential target gene in the monocyte-to-macrophage transition upon exposure to CSF-1[36]. Common to these three DMRs is a potential involvement in monocyte biology specific to a stimulus response. This concept of “priming” for subsequent response to stimulus has been observed with respect to both monocyte activity in CVD [37] and DNA methylation in general [38].

Based on the module- and region-level replication in FHS, we further explored the characteristics of the brown and blue modules. Enrichment analyses of gene-based and locus-based annotations demonstrated that these two modules occupy distinct biological niches. Broadly, the brown module (consisting of about 1000 CpG sites) is enriched for enhancers and other non-proximal regions near immune-related genes, while the blue module (a notably large module of almost 30,000 CpG sites) is enriched for CpG islands near the TSS of development-related genes. One could speculate that these modules also represent different mechanisms of cardiovascular risk: one related to inflammatory burden and the other to long-term effects of early-life exposures, both of which are well-established as contributing to cardiovascular risk [29,39]. Analyses based on cross-tissue epigenome annotations added an additional dimension to these insights by suggesting differential importance of blood cell sub-types for these modules. A cell type specificity analysis, adapted from the eFORGE algorithm [40], revealed the enrichment of monocyte-specific regions of open chromatin (DNase hypersensitivity sites and H3K4me1 peaks) in the brown module. This observation reinforces the idea of monocyte-specific activity suggested by the replicated DMRs as well as that of “monocyte priming” [37]. Based on the tendency of blue module CpGs to be proximal to gene TSS, we focused on enrichment for a promoter-associated marker, H3K4me3, and found a distinct signal related to hematopoietic stem cells. This finding supports a potential mechanisms linking early-life exposure to consequences in adult life [41,42]. We also observed that the blue module was strongly enriched for components of a popular epigenetic age marker [22] as well as for binding regions of the Polycomb-group member SUZ12. As Polycomb-group targets are known to be related to developmental processes [25], this finding contributes additional support to the module’s role as a bridge between development, aging, and disease risk.

It is not clear whether these methylation modules associate with cardiovascular risk upstream, downstream, or independently of traditional cardiovascular risk factors (including age, blood pressure, BMI, smoking, and lipid levels). To explore these relationships, we began by calculating correlations between risk factor levels and blue and brown module activations. Blue correlations were largely weak, while brown correlations were somewhat stronger, following the hypothesis that the blue module is more relevant to early-life, rather than adult, exposures as compared to brown. However, as a semi-stable biological quantity, methylation may have the ability to act as a “molecular recorder” of past exposures, ranging from heavy metals to stress [43,44]. We thus retrieved risk factor measurements from seven prior exams in FHS to compare “cumulative” (calculated as the mean of past exam values) versus current correlations with brown activation. Surprisingly, we observed stronger correlations with cumulative values across almost all risk factors. To address the possibility of confounding in these relationships, we tested linear models predicting brown eigenCpG values from current or cumulative risk factors adjusting for the full set of EWAS covariates. Here, we again observed multiple instances of stronger cumulative relationships, especially for BMI, hsCRP, and triglycerides. Though such a finding could be partially explained by the greater stability in a mean over seven values compared to one, we note that we did not observe this same pattern with respect to the blue module, where associations with current risk factors tended to be stronger. Our observation agrees with a conceptual model in which known risk factors, such as the three noted here, act partially through their cumulative impact over time on immune cell DNA methylation and thus inflammatory processes known to be related to CVD pathogenesis. To more directly test this proposal, we used a basic mediation approach in which we sequentially tested the relationships between cumulative risk factor levels, brown eigenCpG values, and both factors together in predicting incident CVD. Though neither factor exerted a strong effect on the relationship of the other, module activation associations were more weakened after adjustment for cumulative risk factors than the converse. Thus, our models replicate previous findings that cumulative risk factor exposure correlates with CVD risk [45] while suggesting that brown methylation module activation may be sensing, rather than mediating, this effect. One concrete example supporting this observation is the DMR near SLC1A5 containing primarily brown CpGs, one of which (cg02711608) was suggested in Mendelian randomization analysis to be causally downstream of blood pressure [32].

A few limitations should be acknowledged in intepreting the results of this study. First, its observational nature made it impossible to clearly determine causality of the relationships between methylation and cardiovascular risk. While the examination of incident CVD reduced concerns about reverse causation, the discovered associations may only be markers of other disease-causing processes (such as cumulative risk factor exposure, as discussed above). Second, assessment of methylation in blood samples prevented the understanding of potentially causal epigenetic effects in other CVD-relevant tissues. Although some studies report promising findings with respect to blood as a proxy tissue [46,47], and although development-related epialleles may persist across tissues, there is a certain gap in our ability to discover non-blood-related epigenetic patterns in this analysis. Finally, longitudinal methylation measurements were not available in these datasets, preventing an analysis of the intra-individual stability of methylation sites and modules that may be predictive of CVD risk.

The modules and regions discovered in this investigation provide insights into the complex relationships between DNA methylation and cardiovascular disease risk. We show that epigenetic modules track with diverse biological sources of CVD risk, ranging from development- to immune-related processes, and may provide a molecular readout of past exposure to cardiovascular risk factors. We further discover specific differentially methylated regions that may be related to monocyte activation in response to biological stimuli. This work opens the door to further investigation of the epigenetic basis of CVD risk as well as the ability of DNA methylation to act as a biomarker of prior exposures that may be important for disease-relevant prognosis and interventions.

## 5 Methods

### 5.1 Study participants and phenotype collection

Data for the discovery set came from a combined case-control and pseudo case-cohort sampling of 2129 women from the Women’s Health Initiative study, a larger prospective cohort beginning in 1993 that included over 160,000 postmenopausal women from across the United States[48]. Included subjects had no self-reported CVD at baseline, and cases were chosen based on incident centrally adjudicated angina, revascularization, or CHD event during follow-up. Inclusion criteria for methylation measurement resulted in an oversampling of African American and Hispanic participants. Blood samples used for measurement of DNA methylation and clinical biochemistry were taken at Exam 1. Data are available in the dbGaP public repository (accession: phs000200.v11.p3; downloaded on September 27, 2017).

Data for the validation set came from a substudy of the Framingham Heart Study that measured DNA methylation in 2726 subjects from the Offspring Cohort. The Framingham Offspring Cohort was originally established in 1971 to follow 5209 children of the original Framingham Heart Study participants and their spouses[49]. Fasting blood samples for both methylation and clinical biochemistry were collected from participants at Exam 8, which took place from 2005-8. Blood samples were also provided for clinical biochemistry measurements in previous exams, constituting the “past exposures” examined here. Data are available in the dbGaP public repository (accession: phs000007.v29.p10; downloaded on September 27, 2017). Adjudicated cardiovascular event data was collected through 2015, and events were defined here as any of: myocardial infarction, angina pectoris, stroke, or death from CHD (Framingham event codes 1-29).

Blood-based biochemical markers (total cholesterol, LDL-C, HDL-C, triglycerides, glucose, hsCRP, and systolic blood pressure) were log10-transformed for all analyses. In addition, median imputation was used to fill missing values for BMI (20 individuals in total), medication use, and smoking status (thus assuming no medication use and no smoking where these values were missing). Diabetes was defined as either use of diabetes medication or a measured fasting blood glucose level of >125 mg/dL. While directly available in WHI, pack-years of smoking was approximated in FHS by multiplying the number of years since starting smoking by the current number of packs per day.

### 5.2 DNA methylation data processing

In both cohorts, DNA methylation data were collected using the Illumina HumanMethylation450 microarray platform [50] and downloaded as raw intensity files. Preprocessing was performed using the *minfi* and *wateRmelon* packages for R [51,52]. As a quality control step, samples were removed if they showed weak overall signal based on visual inspection of an intensity plot, if they had more than 10% of probes undetected at a detection threshold of p<1e-16, or if the reported sex did not match the predicted sex based on methylation patterns. Probes were removed if they met any of the following criteria: more than 10% of samples undetected at a detection threshold of p<1e-16, location in the X or Y chromosomes, non-CpG probes, cross-hybridizing probes, probes measuring SNPs, and probes with an annotated SNP at the CpG site or in the single-base extension region. Samples were normalized using the Noob method for background correction and dye-bias normalization, followed by the BMIQ method for probe type correction [53,54]. For each dataset, principal components analysis was performed on the set of control probes using code adapted from the CPACOR method of Lehne et al. to account for technical variation [55]. Blood cell counts for 6 blood cell types (CD4+ T-cells, CD8+ T-cells, B-cells, natural killer cells, monocytes, and granulocytes) were estimated using a common reference-based method [56]. After quality control and filtering steps, 422952 (WHI) and 425326 (FHS) CpG sites remained for downstream analysis, formatted as beta values (ratio of methylated signal to total microarray signal). The vast majority of these sites (422688) were available in both datasets.

### 5.3 Weighted gene correlation network analysis

Weighted gene correlation network analysis (WGCNA) was used to find highly correlated modules of CpG sites [17]. The full set of 422952 CpGs passing quality control from WHI were used as input. For computational tractability, blockwise module detection was performed, which treats blocks of features separately for network creation and module detection, followed by eventual merging of highly similar modules. To allow for reasonable computation time, the initial pre-clustering analysis (used to inform the choice of blocks) was performed in a random subset of 100 subjects. A block size of 20,000 was used, and a soft-thresholding power of 8 was chosen to balance approximately scale-free network properties with network connectivity. Unsigned networks were used, based on the fact that the biological consequences of an increase vs. decrease in DNA methylation are much less clear than those of gene transcripts. Whole-module behavior was assessed using the first component from a principal components analysis, performed separately for each module. Scree plots were used to inform the variance explained by each module as well as to justify the use of a single eigenvector as a proxy for module behavior. Module preservation assessment was completed in FHS to confirm cross-dataset robustness of modules. The *modulePreservation* function calculates permutation-based *Z_summary_* statistics reflecting the preservation of density (of within-module adjacencies) and connectivity (maintenance of within-module node correlations) when modules are evaluated in a test set [20]. EigenCpGs were then calculated (according to the principal component weights from WHI), followed by assessment of associations with incident CVD.

Module associations with cardiovascular disease were assessed using Cox proportional hazards regressions, with eigenCpGs as the independent variable and time-to-event measures for incident CVD as the dependent variable. Minimal models adjusted for estimated blood cell counts as well as technical covariates (DNA pull batch in WHI; analysis center + 7 control-probe principal components in FHS – see EWAS section for details). Fully-adjusted models adjusted additionally for biological covariates (age, BMI, smoking status, and pack-years of smoking; sex in FHS; race in WHI). Proportional hazards checks were implemented (cox.zph function in R), and no violations of the Cox regression assumptions were detected at p < 0.05 for any of the modules in WHI or FHS. Mixed models to account for family structure in FHS were also explored, but were found to generate highly similar results (Supp. Table S1).

### 5.4 Epigenome-wide associations of DNA methylation with incident CVD events

For the EWAS analysis, each CpG site was assessed using the the same regression framework as in the module-based models, separately in both WHI and FHS. Methylation beta-values replaced eigenCpGs as the independent variable, and the full set of technical and biological covariates was used. To remove the influence of beta-value outliers, samples were excluded for each CpG if their beta value was outside of the interval [25%ile - 3 * *IQR*, > 75%ile + 3 * *IQR*]. QQ plots and calculation of the genomic inflation factor λ revealed that genomic inflation was not initially adequately controlled in FHS, but after additional adjustment for 7 CPACOR principal components (chosen based on a Scree plot assessment of CPACOR results), a reasonable inflation of λ = 1.09 was achieved. CPACOR uses principal components analysis on the set of control probes from the methylation array in order to estimate and control for potential batch effects without disturbing biological signal [55]. Proportional hazards checks were implemented as in the module-based analysis for the top EWAS hits in WHI, and no systematic departure from the Cox regression assumptions were detected.

Comb-p, implemented as a Python module, was used to call differentially methylated regions (DMRs). The algorithm takes as input p-values from the EWAS, removing the requirement for additional covariate adjustment. Comb-p first calculates an autocorrelation function (ACF), for which a maximum distance of 1kb and a step size of 50 bases were used. Next, it uses the ACF to adjust each p-value using a Stouffer-Liptak-Kechris correction, followed by identification of contiguous regions of sites with adjusted p-values below some threshold (here, p<0.1 with no more than 500 bases between neighboring sites in a region). Finally, the ACF is recalculated out to the maximum region size (a step size of 50 was used here as well) and regional p-values are calculated using the Stouffer-Liptak test. For multiple testing correction of DMRs, Comb-p calculates the number of effective tests separately for each DMR as the number of loci tested divided by the size of the region (typically ~1kb).

DMRs were examined to evaluate whether their constituent CpGs contained any residual SNPs-under-probe that escaped filtering based on the Illumina HumanMethylation450 annotation. These checks were performed manually using the UCSC Genome Browser [57] and a dbSNP-based annotation track displaying common (>=1% minor allele frequency) variants.

### 5.5 Module enrichment analyses

Gene ontology-based enrichment analysis of modules was performed using the gometh function from the *missMethyl* package for R [58]. In this procedure, CpG sites are annotated to genes using the Human-Methylation450 microarray annotation from Illumina, resulting in a binary vector indicating whether each gene is associated with any of the CpG sites of interest (for example, CpGs constituting a module). Prior probabilities for each gene being selected are estimated based on the total number of associated CpG sites on the array. Enrichment analysis is then performed for each gene ontology category using Wallenius’ noncentral hypergeometric distribution, which generalizes the basic hypergeometric distribution to account for biased sampling.

Locus-based enrichment analyses were performed using basic two-tailed hypergeometric tests for overlap between module membership and annotation category membership. CpG annotations with respect to both CpG islands (Island, North shore, Open sea, etc.) and genes (TSS1500, 3’ UTR, Body, etc.) were retrieved from the standard Illumina HumanMethylation450 microarray annotation. CpG sites were annotated for Polycomb-group target status using embryonic stem cell SUZ12 binding regions retrieved from Lee et al. [26].

### 5.6 Inference of cell type specificity

Epigenomic annotations were used to test for relative enrichment of module CpGs in cell type-specific regulatory regions. Annotations for broad peaks in DNase sensitivity as well as ChIP-seq signal for H3K4me1 and H3K4me3 were obtained for 6 blood cell types (monocytes, natural killer cells, T-cells, B-cells, and hematopoietic stems cells from males and females) from the NIH Roadmap Epigenomics Project database [21]. For each combination of epigenomic feature and cell type, CpGs from the HumanMethylation450 array were classified as to their membership in a peak region. Relative enrichments of in-peak CpGs for modules were then calculated as the ratio of 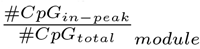 to 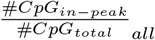 and presented as *log*_2_(relative enrichment) for ease of visualization. Cell type specificity of different modules can then be compared by examining relative enrichments across cell types, especially with respect to highly-represented regulatory annotation types (e.g. DNase hypersensitivity sites for a module enriched in enhancers). We note that this method borrows from the permutation-based eFORGE tool methodology [40], which could not be used here due to the size of the blue module. However, we confirmed similarity of our results to those from the eFORGE method for the brown module (Supp. Fig. S3).

### 5.7 Risk factor integration

Risk factors were incorporated into the module-based analysis in a series of steps. First, Pearson correlations between risk factor levels and module eigenCpGs were calculated to provide a high-level understanding of the strength of their relationship. Risk factors in WHI were all measured at Exam 1 (concurrently with the methylation measurement), while risk factors in FHS were collected for all exams prior to and including Exam 8 (the time of the methylation measurement). In FHS, correlations with past risk factor levels as well as a “cumulative” exposure level (equal to the mean of each set of risk factor levels from Exam 1-7) were also calculated.

Next, linear models were used to assess these same module-risk factor correlations in FHS while adjusting for potential confounding variables. These models predicted module eigenCpGs using either cumulative (Exams 1-7) or current (Exam 8) risk factors, while adjusting for the same set of technical and biological covariates as in the EWAS (described above). In this step, both eigenCpGs and risk factors were standardized before modeling in order to facilitate effect size comparisons across risk factors and across modules.

Finally, the relationship between cumulative risk factors, the brown module, and incident CVD was examined, using the same Cox regression setup as in the EWAS to perform a basic mediation analysis for BMI, hsCRP, and triglycerides. Here, cumulative risk factor exposure (as defined above) acted as the exposure, brown methylation module activation (represented by the brown eigenCpG) acted as the mediator, and incident CVD acted as the outcome. Having established the strong exposure-mediator links, three subsequent Cox models were examined: cumulative risk factors only, brown eigenCpG only, and both simultaneously. All models adjusted for the full set of technical and biological covariates as well as the “current” level (i.e. at Exam 8) of the risk factor in question.

## Supporting information

