## Supplementary material for "DNA methylation modules associate with incident cardiovascular disease and cumulative risk factor exposure"

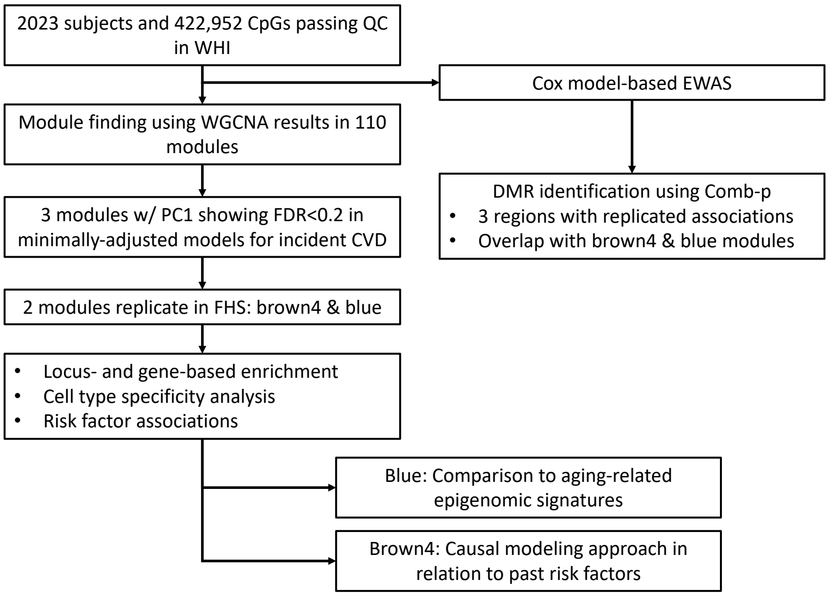


Figure S1: Study overview, including module- and region-based analyses as well as follow-up.

Table S1: P-values for module associations with incident CVD in discovery and replication.

| | WHI (discovery) | | FHS (replication) | | |
|---|---|---|---|---|---|
| Module | Partially adjusted | Fully adjusted | Partially adj. | Partially adj. (mixed) | Fully adjusted |
| blue | 0.0002736 | 0.0500018 | 0.0000085 | 0.0000085 | 0.8189348 |
| brown4 | 0.0045462 | 0.0872688 | 0.0000962 | 0.0000963 | 0.0997390 |
| lavenderblush3 | 0.0050028 | 0.0210976 | 0.0202819 | 0.0202804 | 0.1580861 |

[1] Partially-adjusted models are adjusted for technical covariates (DNA pull batch in WHI and study center + 7 control probe PCs in FHS) and estimated cell counts. Fully-adjusted models are additionally adjusted for age, sex, smoking status and smoking pack-years.
[2] Mixed model contains a random intercept for each family.

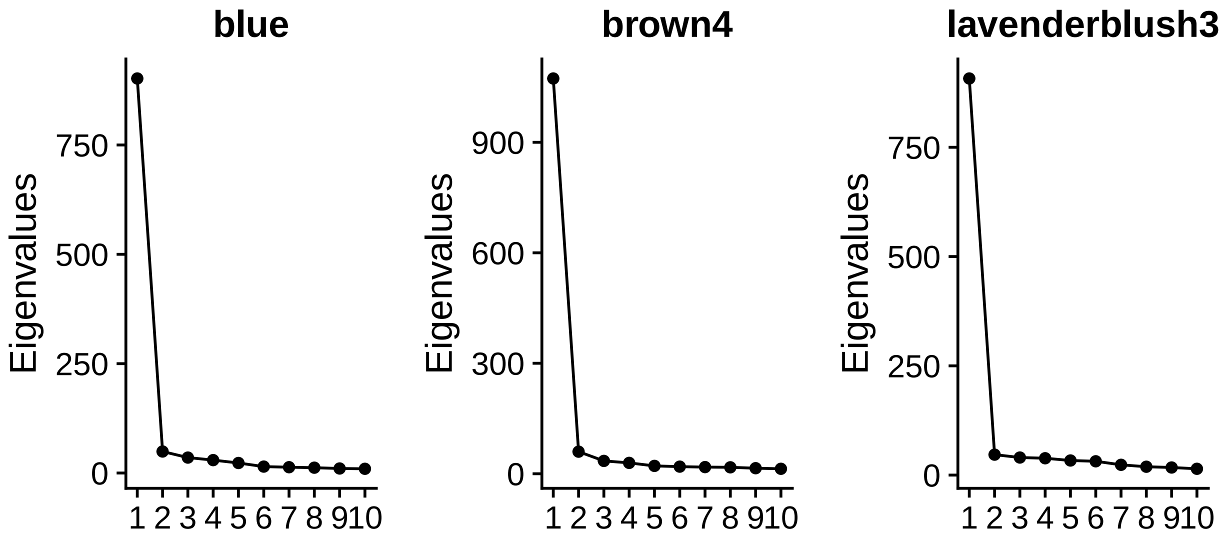


Figure S2: Scree plots for PCA on the set of CpGs corresponding to each of the top modules.

Table S2: CpGs with FDR < 0.05 in the discovery set (Bonferroni threshold = 1.18e-7)

| CpG | Chromosome | Dir. of Assoc. | P-value | Location | Annotated Gene | Replication P-value |
|---|---|---|---|---|---|---|
| cg09155044 | chr16 | + | 6.63e-09 | TSS1500 | VKORC1 | 0.107 |
| cg24434800 | chr1 | + | 5.04e-08 | | | 0.629 |
| cg11691298 | chr2 | + | 1.1e-07 | Body | FAM59B | 0.525 |
| cg02379107 | chr20 | + | 4.72e-07 | TSS1500 | KIAA1755 | 0.930 |

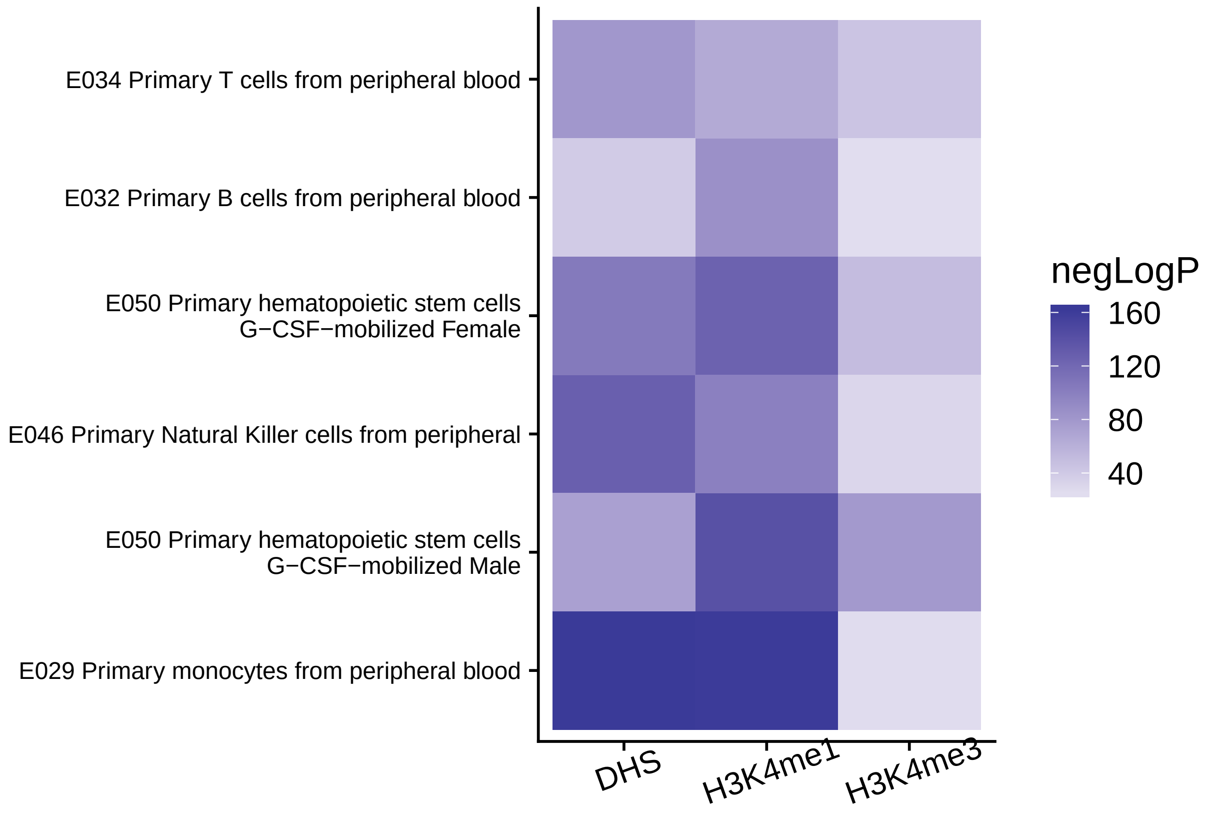


Figure S3: eFORGE cell type-specificity plot for the brown4 module.

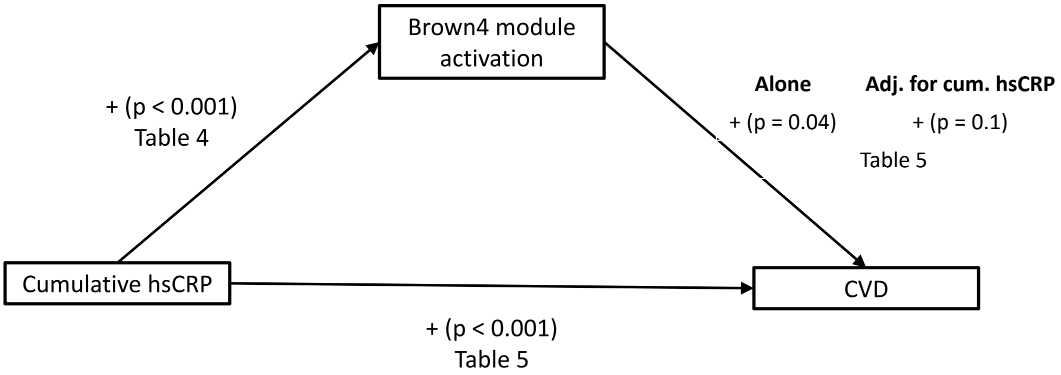


Figure S4: Example diagram of cumulative risk factor mediation by brown4 methylation module activation. Results from 4 regressions are shown: cumulative risk factor exposure to brown4 activation, cumulative risk factor exposure to incident CVD, and brown4 activation to incident CVD with and without adjustment for cumulative risk factor exposure. Regression terms represented as: sign of coefficient (p-value).
